# A spatialised agent-based model of NOTCH signalling pathway in Endothelial Cells predicts emergent heterogeneity due to continual dynamic phenotypic adjustments

**DOI:** 10.1101/2022.08.06.503043

**Authors:** Francois Chesnais, Timothy J Sego, Elena Engstler, Matteo Battilocchi, Davide Danovi, James A Glazier, Lorenzo Veschini

## Abstract

Vascular Endothelial Cells (EC) plasticity is key to homeostasis and its disruption is a hallmark of diseases such as cancer, atherosclerosis, and diabetes. The EC lineage has evolved to address in parallel sensor and actuator functions. This ability is reflected in remarkable phenotypical heterogeneity of EC across different tissues, within the same tissue, and within the same vascular bed as demonstrated by single cell image analysis and transcriptomics studies. However, how the molecular signalling dynamics in EC could generate and maintain such heterogeneity in different contexts is still largely unexplored. Recently we reported that confluent EC have spatially heterogeneous NOTCH signalling pathway (NSP) levels in vitro as confirmed from analysis of available OMICS databases. Here, we show that spatial heterogeneity of NSP levels is a feature of aortic murine endothelia *in vivo* and recapitulated by human EC in culture despite absence of signalling from mural cells. We study lateral induction and inhibition, cis-interactions and signalling, and target genes autoregulation in NSP. Using mathematical models and experimental observations we report that NSP dynamics can generate stable, periodic, and asynchronous oscillations of the NSP target HES1. Importantly, we observe that cell contact dependent NSP signal oscillations is the most likely parsimonious mechanistic hypothesis justifying observed spatial heterogeneity in endothelia. We propose that NSP is sufficient to enable individual EC in monolayers to acquire different phenotypes dynamically explaining robustness of quiescent endothelia in performing parallel functions.

## Introduction

Endothelial cells (EC) compose the inner lining of all blood and lymphatic vessels in the human body and they are phenotypically heterogeneous across different tissues. EC phenotypic heterogeneity is an evolutionarily conserved feature emerging early during embryonic development (Aird, 2007a, 2007b). Disruption of EC homeostasis is a recognised hallmark of diseases such as cancer and atherosclerosis and EC are primary targets for pharmacological interventions due to their key role at the interface between blood and all tissues (Potente et al., 2011; Simmons et al., 2005). Previous *in vivo* studies (Lee et al., 2022; McCarron et al., 2019) have shown that EC in large vessels have heterogeneous phenotypes (e.g., differential Ca++ sensitivity). We have previously demonstrated the value of measuring images of cultured EC to assess phenotypical and intracellular signalling heterogeneity in endothelial monolayers using our EC profiling tool (ECPT)(Chesnais et al., 2022). Our work using ECPT has established that quiescent EC within the same monolayer *in vitro* display heterogeneous levels of NICD (second messenger of NOTCH Signalling Pathway, NSP) and HES1 (NSP target gene), echoing results obtained in cancer cells (Sabherwal et al., 2021).

Establishing mechanistic links between signalling pathways such as NOTCH and heterogeneous EC phenotypes is key to improve our understanding of EC biology, to develop more effective drugs and increase predictability of treatments outcome (Zhou et al., 2022). For example NSP has been implicated in both atherosclerotic plaques formation and regression (Kong et al., 2022; Vieceli Dalla Sega et al., 2019).

Previous work *in vivo, in vitro* and *in silico* has defined a framework to elucidate emerging NOTCH-dependent phenotypes. The current hypothesis is that mechanisms of lateral induction (LId) and lateral inhibition (LIb, trans-interactions) as well as ligand-receptors interactions in the same cell (cis-interactions, CI) (Boareto et al., 2016; Chesnais et al., 2022) coexist to induce differential signalling levels (e.g., levels of HES1) and phenotypes. Beyond the well-known stable alternating spatial patterns in NSP observed in several contexts (e.g., developing mouse retina neuroepithelium) (Formosa-Jordan et al., 2013), temporal oscillations in NSP (levels of NICD or NSP target genes) have been predicted *in silico* and observed experimentally (Marinopoulou et al., 2021; Sabherwal et al., 2021; Ubezio et al., 2016) The current mechanistic hypothesis is that NSP levels oscillations may result from cell autonomous negative autoregulation of HES1 transcription (Hirata et al., 2002). HES1 is a well-established NSP target gene in EC which differential expression regulates functions such as cell proliferation and inter-cellular junctional stability (Bentley et al., 2014; Fang et al., 2017; Fernández-Martín et al., 2012). In several biological contexts, variations in the levels of NSP in neighbouring cells determines differential cell fates and EC constitutively express NOTCH ligands and receptors independently from NSP(Curry et al., 2006). NSP is key to tip cell selection and tip-stalk cell crosstalk during VEGF induced neo-angiogenesis (Jakobsson et al., 2010, 2009). However, the spatio-temporal dynamics of NOTCH signalling in “stable” (non-proliferative, non-migratory) EC monolayers (i.e., endothelia) is poorly understood. In mathematical formalisation, a negative autoregulatory loop can only oscillate in the presence of significant delays in regulation; else, negative autoregulation leads to stable graded expression. Delays could result from transcription factor’s nuclear import dynamics, transcriptional/translational delay or the inclusion of additional intermediate factors in the signalling chain. However, HES1 appears to act as a simple direct auto-inhibitor without intermediate factors.

Our previous results show that HES1 is repressed in few cells within a confluent EC monolayer suggesting that these cells could be licensed for proliferation (Chesnais et al., 2022). Furthermore, these results suggest a non-stable NSP dynamics in EC monolayers paralleling results from angiogenic EC (Ubezio et al., 2016) and breast cancer cell lines (Sabherwal et al., 2021).

Directly measuring HES1 dynamics in primary EC to verify this hypothesis is currently challenging. However, multi scale models calibrated on experimental data enable a qualitative evaluation of hypotheses on the dynamic of cellular systems signalling when direct experimental measures are inaccessible.

Here, to establish a mechanistic link between NSP levels and acquisition of differential phenotypes in EC we firstly aimed to confirm that spatial heterogeneity of NSP is also appreciable *in vivo*. Then we aimed to evaluate *in silico* whether NSP alone is in principle sufficient to generate such heterogeneity in space and time. We used time course experiment where we inhibited NSP in human umbilical vein and human aortic EC (HUVEC and HAOEC) and measured time variations in nuclear NOTCH1 and HES1 at single cell and population levels by ECPT. We used experimental cell maps and corresponding data to calibrate a novel spatialised multi scale model (SMSM) of NSP encompassing LIb, LId, CI, and HES1 autoregulation. Finally, we used our SMSM to evaluate whether and under which conditions synthetic data reproduce the spatial distributions of NOCTH signal observed *in vitro* and *in vivo*.

## Results

### Nuclear HES1 have heterogeneous intensities in murine aortic EC

We first set to address whether NSP levels heterogeneity (NSH) we previously measured *in vitro* is a feature of endothelia *in vivo* where NSP might be affected by co-signalling from mural cells (Baeten and Lilly, 2017). To evaluate whether endothelia in vivo demonstrated spatial NSH we performed en face stainings of murine aortas as described before (Hakanpaa et al., 2015). Fig.1A shows a representative image of murine aortic endothelium. HES1 immunostaining (yellow) shows heterogeneity in intensity of signal especially, the presence of few EC with low- or high-signal like observed *in vitro* (Fig 1C). Fig. 1B (green trace) shows single-cell quantification of HES1 intensities across 3 images taken from 3 independent samples confirming heterogeneous distribution of HES1 intensities across aortic monolayers. Black and red overlay traces in Fig. 1B and maps in Fig. 1C correspond to measures in HAoEC and HUVEC monolayers demonstrating that *in vitro* data qualitatively resemble *in vivo* scenario.

**Fig 1:**
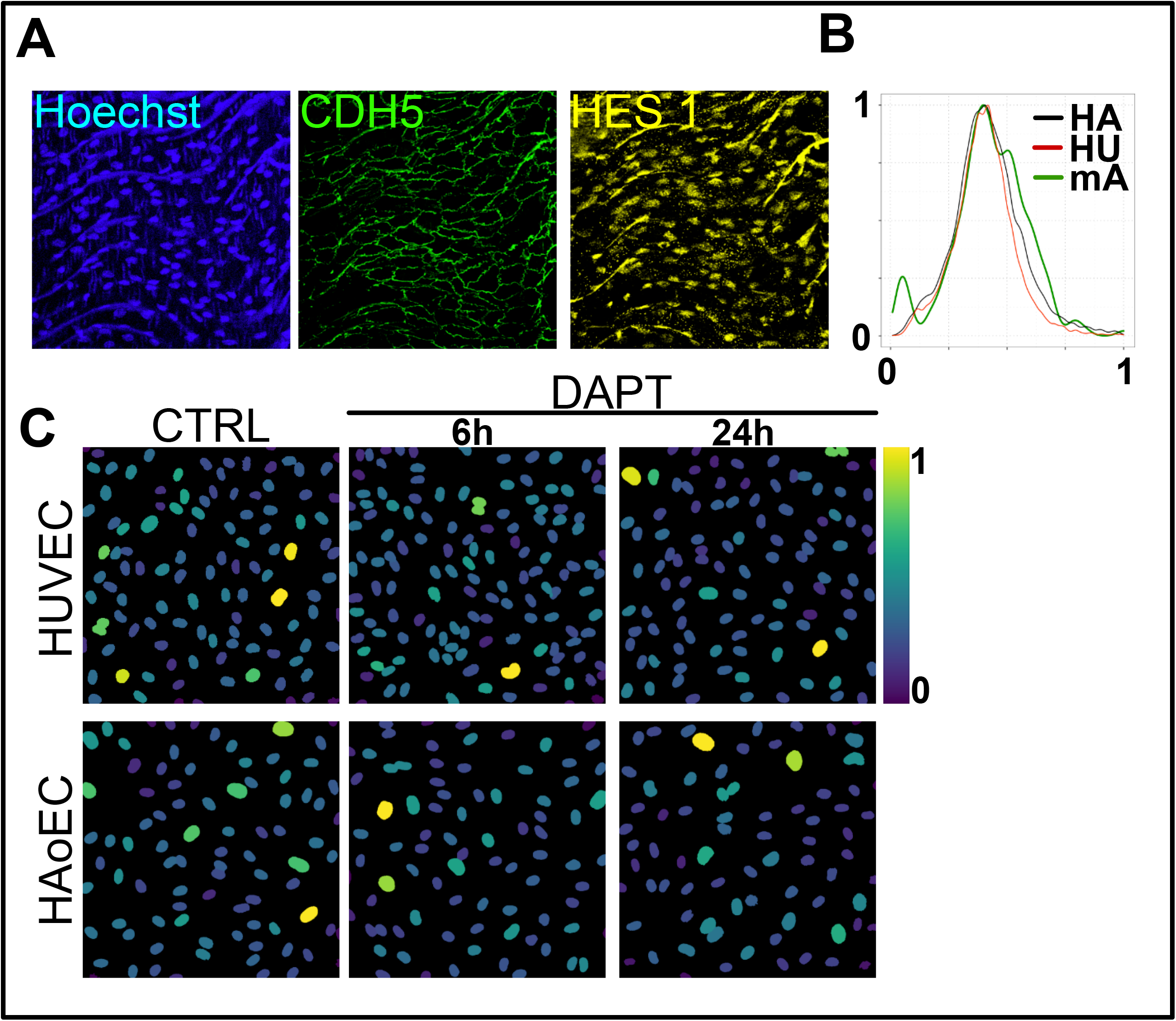
HES1 is spatially heterogeneous in endothelia. **A)** Confocal images of murine aortic endothelium immunostained for CDH5 (green) and HES1(Yellow). DNA was counterstained with Hoechst (Blue). Each image corresponds to maximal projection of a 20μm confocal stack. **B)** Density distribution plot of normalised HES1 intensities corresponding to mEC in aortic endothelium (green trace), HUVEC (red trace) and HAoEC (black trace). **C)** Representative HES1 maps obtained by ECPT analysis of HUVEC and HAoEC at baseline or treated with DAPT 5μM for 6 or 24 hours. Colour scale represents normalised HES1 intensities.

Overall, these results demonstrate NSH in murine aortic endothelia and that the mechanisms underpinning observed heterogeneity are preserved *in vitro* despite absence of signalling from mural cells. These results strongly suggest that the molecular mechanisms driving NSH are predominantly intrinsic to EC validating EC cultures as models to investigate them in a controlled experimental setup.

To investigate the underlying molecular dynamics driving NSH in endothelia we developed a computational framework encompassing established and hypothesised rules driving NSP dynamics. We calibrated our framework with experimental data and cell maps from standardised *in vitro* EC cultures and we explored which hypotheses could reproduce experimental data *in silico.*

### Creation and calibration of a spatialised multiscale model of NOTCH signalling in EC

Assuming that NOTCH signalling is sufficient to generate the heterogeneous maps we observed *in vitro* and *in vivo,* then NSH could emerge either due to generation of multiple stable phenotypes or alternatively instable, oscillating ones as reported previously (Hirata et al., 2002; Marinopoulou et al., 2021; Sabherwal et al., 2021; Shimojo et al., 2008; Yoshioka-Kobayashi et al., 2020). To evaluate these hypotheses in the context of EC monolayers we built a spatialised multiscale cellular model of NSP including Lateral Inhibition (LIb) and Lateral Induction (LId) as described in previous work (Boareto et al., 2016; Sprinzak et al., 2010). We based our sub cellular Proteins/Gene Regulatory Network (P/GRN) model on previous frameworks (Boareto et al., 2015) following similar deterministic formalism and expanded it to include productive cis interactions (CI) (Nandagopal et al., 2019) (Fig. 2A) and HES1 autoregulation (Monk, 2003). We started assigning previously established values to all model parameters (e.g., production degradation rates of receptors, ligands second messengers and TF). We embedded the P/GRN model in a spatialised context using either regular cell dispositions (square lattice of identical cells) or experimentally derived cell maps (Fig 2B).

**Fig 2:**
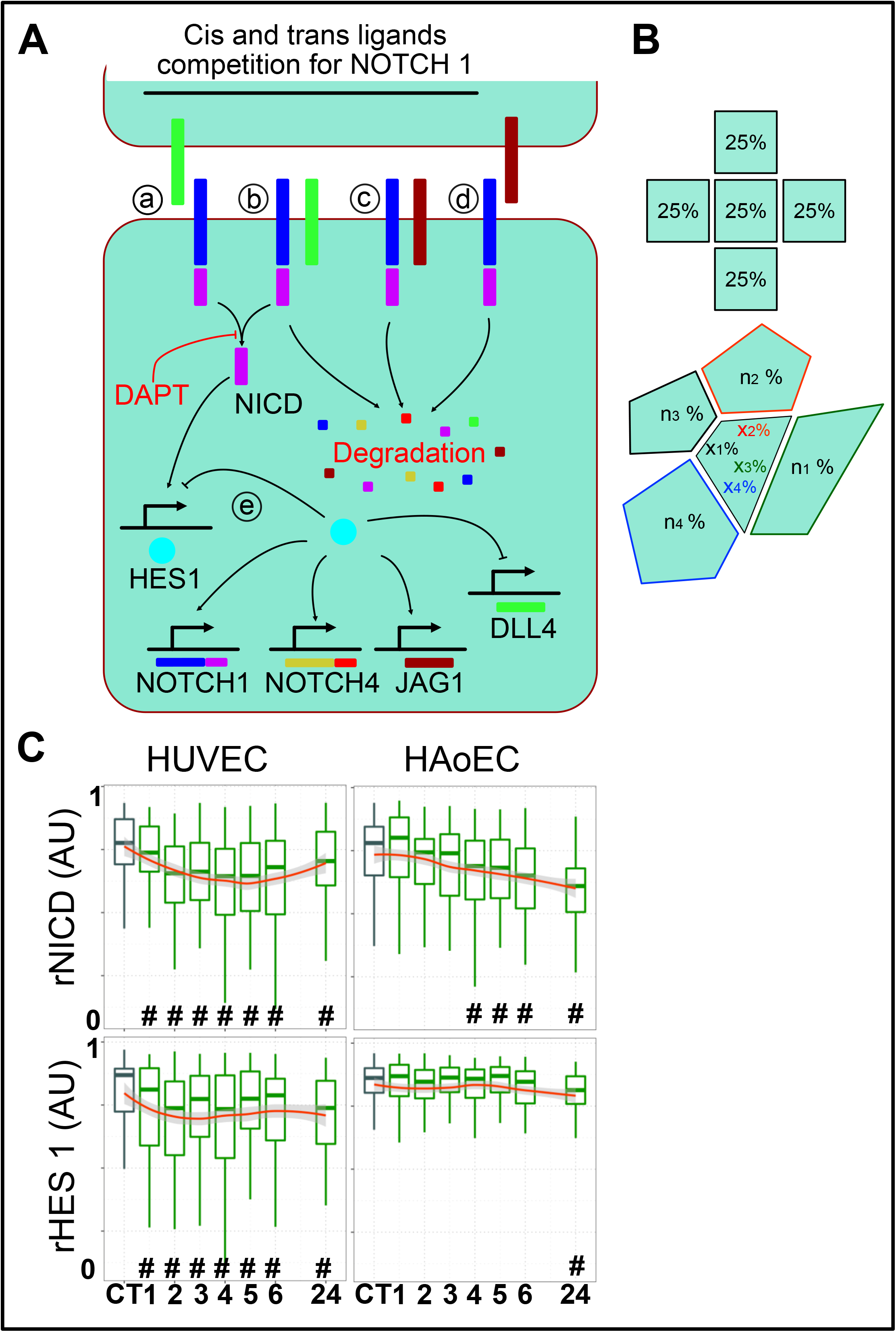
Multiscale model of NOTCH signalling in EC monolayers. **A)** Schematic depicting the molecular interactions captured in our SMSM. a-productive Trans Interaction (TI) of Dll4-NOTCH1, b-productive/non-productive Cis Interaction of Dll4-NOTCH1, c-non-productive CI Jag1-NOTCH1, d-non-productive TI Jag1-NOTCH1, e-HES1 autoregulation. **B)** Schematic representing interactions between regularly disposed square cells compared to irregular disposition of irregular polygonal cells. In the former, total available ligand/receptors are shared equally with neighbours. In the latter, differential membrane contact area leads to different extent of shared ligands/receptors with neighbours. X_1-4_% amounts of ligands/receptors shared by the central cell (i.e., a fraction proportional to shared area with different neighbours). N_1-4_% amounts of ligand/receptors shared with central cell (i.e., a fraction dependent on each neighbour’s interactions with its own other neighbours). **C)** Box plots of multiple (individual images) KS distance measures from reference ECDF (Untreated cells, grey) for HUVEC and HAoEC, either untreated or treated with DAPT for the indicated number of hours (green boxes). # p<0.01 against control (CT, untreated cells, grey boxes).

The timescale of NSP transduction can be affected by contextual/phenotype-dependent factors. To further specify our P/GRN model to EC we experimentally timed degradation of NICD and HES1 in cultured EC. To this aim, we timed the variations in nuclear NOTCH and HES1 signal in EC upon exposure to the gamma secretase inhibitor DAPT. DAPT limits NOTCH signal transduction by inhibiting cleavage of internalised NOTCH receptors to produce the second messenger NICD (Fig. 2A). We measured nuclear NOTCH1 and HES1 in EC monolayers exposed to DAPT for different times using the ECPT (Fig. 2C)(Chesnais et al., 2022). Fig. 2C shows normalised plots for relative NICD and HES1 intensities (1/KS-distance from cumulative control condition) in HUVEC and HAoEC populations upon treatment with DAPT. These results demonstrate that half maximal NICD inhibition in our experimental conditions is achieved in 1 hour in HUVEC and in 4 hours in HAoEC which is coherent with higher intrinsic signalling in the latter (Chesnais et al., 2022; Luo et al., 2020). Long term DAPT treatment (24h) resulted in further NICD inhibition in HAoEC but not in HUVEC (Fig 2C). Overall, these results demonstrate that cellular NICD degrades at fast rate (minutes) in line with previous models and data (Boareto et al., 2015; Hirata et al., 2002; Marinopoulou et al., 2021). Furthermore, our results highlight that the net effects of NSP perturbation are unsurprisingly cell phenotype specific. Similarly, we estimated the time scale of HES1 inhibition in the same cells. The plot in Fig 2C shows that half maximal inhibition of HES1 in HUVEC is achieved in one hour paralleling NICD results. However, HES1 levels in HAoEC were only affected at later times >6 hours upon DAPT exposure. Overall, results in HUVEC closely mirror previous results using different experimental systems suggesting that the underlying NOCTH dynamics investigated in previous works is paralleled in EC (Boareto et al., 2015; Monk, 2003; Nandagopal et al., 2019; Yoshioka-Kobayashi et al., 2020). Using these results and reference parameters defined in previous work we set the granularity of our simulation to 5 minutes/MCS and manually calibrated NICD and HES1 production/degradation rates in our model to match these results. Our estimates largely reflect previously established results suggesting that the core NOTCH molecular machinery has similar characteristic timescale in EC as in other cell types (Boareto et al., 2015; Monk, 2003).

### In silico analysis of NOTCH signalling in EC monolayers reveals highly dynamic scenarios

To evaluate the qualitative responses of our calibrated model against selected parameters and to validate it against previous models, we performed a coarse-grained parameter scan. We started from regular cell dispositions and subsequently expanding the results to experimental cell maps (SFig. 1 and Fig. 3A). Analysis of parameter scan data revealed that our model can generate several qualitatively different scenarios under different parameter settings. Fig. 3A shows density distributions corresponding to synthetic data for HES1 pooled from 10 independent runs under indicated parameters (bD4, bN1 and Vmax J1 and N4 all set to 1) demonstrating a strong influence of the parameter cr (regulating the strength of CI) on the qualitative output of the model. Plots in Fig. 3A are derived from runs employing experimental cell maps however very similar results were obtained with regular cell dispositions (SFig. 1). Fig. 3A shows snapshots of cell maps and time tracking data corresponding to five possible scenarios occurring under different parameter settings as indicated (in represented scenarios all parameters are set to default values, 0 for *OFF* state or 1 for *ON* state except *cr and delay*). In absence of LId mechanism (no production of Jagged 1) the model predicts generation of stable checkerboard patterns with alternating high/low phenotypes (Fig. 3Bi). Furthermore, differently from previous models and our own data using regular cell dispositions (Boareto et al., 2015) and SFig. 1), we could observe emergence of intermediate phenotypes due to differential contact between cells. Introduction of LId (production of Jagged 1) and moderate CI (*cr*< 0.05) favoured the emergence of stable intermediate phenotypes (Fig. 3A ii, iii) similarly to regular cell dispositions (SFig. 1) and in line with previous data (Boareto et al., 2015). Finally, introduction of sustained CI (*cr* = 0.05-0.1) caused cells to oscillate between high and low phenotypes (Fig. 3B) similarly to regular cell dispositions (SFig. 1). The oscillations were periodic, and their character period and amplitude were sensitive to variation in the simulation parameters we considered especially *cr* and *delay* in HES1 processing. Taken together our results qualitatively reproduce previously described behaviours and extend the analysis to non-regular cell dispositions. Our data show that introduction of asymmetric signalling between neighbouring cells can produce intermediate phenotypes independently from LId and CI mechanisms. Furthermore, our data show for the first time that sustained CI (extent of CI ~ extent of TI) can in principle produce oscillatory phenotypes in a cell monolayer with either regular or irregular cell dispositions (SFig. 1 and Fig. 3Biv). In a purely LIb mechanism without CI, the checkboard pattern emerges due to asymmetry in Dll4 expression in neighbouring cells which is stable because repressed receiver cells cannot signal through Dll4. Introducing CI can break this symmetry because signalling sender cells can auto-repress DLL4 via CI becoming an intermediate sender-receiver cells (SFig. 2). We initially hypothesized that oscillations could also in principle emerge due to an autoregulatory feedback loop with delay in the TF HES1 as previously reported (Hirata et al., 2002) and included this aspect as described in methods. However, our simulations failed to reproduce robust HES1 oscillation by cell-autonomous autoregulatory feedback using previously described parameters and instead produced short-lived oscillations which dampened after few MCS (Fig. 3Av).

**Fig 3:**
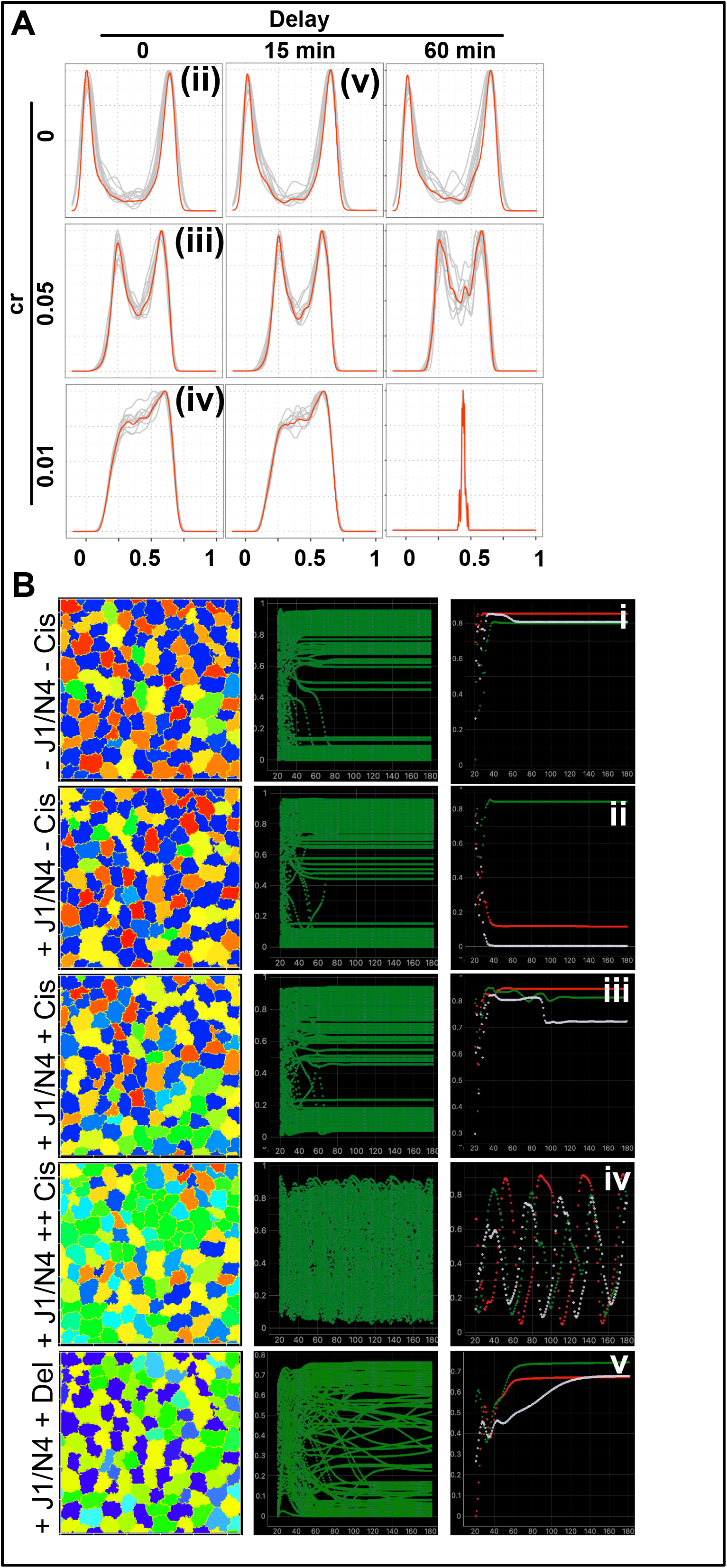
In silico analysis of NOTCH signalling in EC monolayers reveals highly dynamic scenarios. **A)** Density distribution plots corresponding to synthetic data under indicated cr and delay parameters (for bD4, bN1. J1=1). Red traces indicate kernel density estimates corresponding to accumulated results from 10 independent simulations. Grey traces indicate kernel density estimates of individual simulations. (ii-v) indicates corresponding representations in B. **B)** Representative maps (left panels) and timeseries (mid panel, all cells or 3 selected cells right panels) for the indicated conditions (−= OFF, += ON).

Overall, our observations *in silico* suggest that in the context of EC (cell monolayers constitutively expressing NSP ligands and receptors) oscillatory behaviours of NSP are in principle possible and likely due to cell-contact dependent mechanisms rather than cell-autonomous TF autoregulation. Furthermore, our results highlight a key role of cell shapes, dispositions, and spatial relationships in producing different qualitative response of NSP.

### Heterogeneous HES1 in monolayers implies oscillatory phenotypes of individual EC

To evaluate whether any of our theoretical prediction scenarios could reproduce experimental data we performed a fine-grained parameter scan over our set of selected parameters and compared each result against experimental data (Table 1, Methods). To facilitate a fine-grained parameter scan while maintaining the total number of simulations reasonably low we implemented a random walk search followed by iterative optimisation as described in methods and SFig. 3. Fig. 4A shows scatterplots of all simulations in our parameter scan demonstrating that under certain parameter sets our model was able to produce results closely resembling experimental data (positive hits hereafter, KS D<0.1, green-yellow dots).

**Fig 4:**
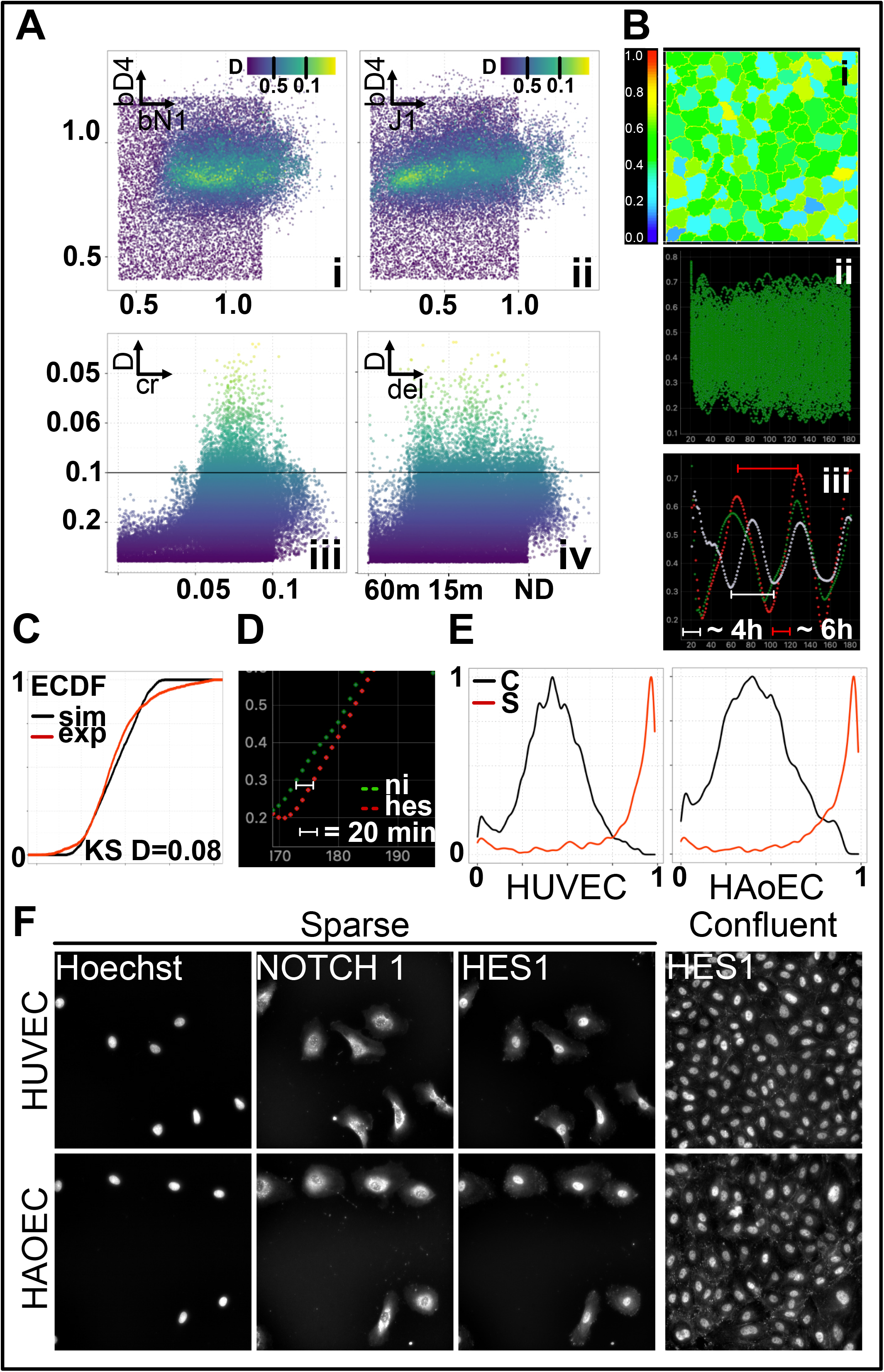
Heterogeneous HES1 in monolayers implies oscillatory phenotypes of individual EC. **A)** Scatter dot plots representing results of a fine-grained parameter scan across bD4, bN1, J1, cr, and delay parameters as indicated in individual plots. Colour scale correspond to KS distance values against reference ECDFs (Experimental data for HUVEC and HAoEC), green/yellow dots KS D<0.06. **B)**. Representative cell map (i) and HES1 time series (all cells, ii and three random cells, iii). Annotations in iii indicate distance between successive peaks in timeseries of corresponding colour and approximate length in hours (1h = 10 MCS). **C)** ECDFs corresponding to representative simulation in B (black trace) compared with experimental data (HUVEC, red trace) and corresponding KS D value. **D)** Zoom in of time series for one random cell for simulation in B showing NICD (ni, green trace) and HES1 (hes, red trace). Delay between onset of NICD and HES1 is approximately 20 minutes (4-5 MCS). **E)** Density distribution estimates of HES1 intensities in confluent (black trace) or sparse (red trace) EC (HUVEC or HAoEC as indicated). **F)** Representative fluorescence images (Nuclei, Hoechst, NOTCH1 and HES1 as indicated) of sparse or confluent EC (HUVEC or HAoEC as indicated).

Fig. 4Ai and ii show that positive hits were restricted in a relatively narrow region of possible bD4, bN1 and J1 values (basal or inducible production rates of Dll4, NOTCH1 and Jag1 respectively) which correspond to physiologic ranges as per model’s specifications (table 1 and methods).

Scatterplot in Fig. 4Aiii demonstrate that positive hits were only possible when *cr* parameter was bigger than ~ 0.05 corresponding to sustained CI and typically producing solution encompassing oscillations in HES1, the best solutions were obtained with *cr* values in the 0.06-0.07 range. Scatterplot in Fig. 4Aiv demonstrate that, in our model, delay in the HES1 autoregulatory feedback doesn’t have a major impact on fitness to experimental data, top solutions (D<0.07) were possible with either unrealistically high (>60 min) or negligible delay values. Fig. 4B shows simulation results corresponding to one representative positive hit scenario with KS D= 0.08 (Fig. 4C). Fig 4B i show a snapshot cell map of HES1 levels at MCS 180 (18h) of the simulation which is qualitatively similar to experimental cell maps in Fig 1C. Fig 4B ii and iii show timeseries plots of HES1 levels of the same simulation and representing traces corresponding to all cell in the simulation (ii) or three randomly selected cells (iii). Annotations in Fig 4B iii highlights period of oscillations of 4-6h which was a typical result among many of the positive hits analysed and matching previously reported experimental data (Ubezio et al., 2016). None of the positive hit scenarios had stable NSP levels suggesting that oscillating NSP levels might be a key feature of quiescent endothelia.

Overall, our results strongly suggest that NSP is in principle sufficient to generate endothelial NSH we observed *in vitro* and *in vivo*. Furthermore, the underlying mechanism generating HES1 spatial heterogeneity in EC monolayers is likely to encompass oscillations in the TF in individual cells as also suggested by previous results (Ubezio et al., 2016). Differently from previous hypotheses which also related to different biological context, our model does not support cell-autonomous mechanisms as origin of HES1 oscillations prompting us to verify this hypothesis experimentally.

### Oscillatory behaviour of HES1 in EC is likely mediated by cell-contact dependent mechanisms

To verify whether HES1 oscillations in EC predicted by our model are cell-contact dependent we measured HES1 expression in sparse EC with minimal EC-EC contact. We hypothesized that heterogeneity produced by cell-contact dependent oscillations should be lost in isolated cells. Instead, if we could observe significant heterogeneity in isolated cells, it would directly imply that oscillations are generated by cell-autonomous mechanisms. Fig. 4E shows plots density distributions of HES1 in sparse cells (S, red traces) compared to confluent cells (C, black traces) and showing that HES1 expression in sparse cells was homogeneously high and low HES1 expressing cells were absent. Fig. 4F shows representative immunofluorescence on sparse and confluent cells demonstrating absence of HES 1 heterogeneity in isolated cells. Notably, NOTCH1 immunostaining in sparse cells displayed active signalling as shown by nuclear NOTCH1 intracellular domain (punctuated peri and intra-nuclear stain) demonstrating signalling in absence of cell contact. These results suggest that spatial heterogeneity in HES1 levels is lost in sparse cells and that potential oscillations are mediated by cell contact. Furthermore, since both nuclear NOTCH1 and HES1 signals were invariably high in sparse cells, as also previously reported (Curry et al., 2006), we can support a hypothesis where productive CI mediates sustained HES1 production. We conclude that oscillations in EC monolayers predicted computationally and supported by experimental evidence are likely cell-contact dependent and, within our computational model, dependent on CI.

## Discussion

EC heterogeneity across tissues, within the same tissue and even within the same vascular bed is a reflection of plasticity and how the EC lineage has evolved to address several different sensor and actuator functions in a highly parallel fashion (McCarron et al., 2019, 2017).

Transcriptomic profiling of EC from different tissues and organs reveals similar NSP levels heterogeneity (NSH) in EC within the same vascular bed (Kalucka et al., 2020). Our recent results show that confluent EC derived from the same vascular bed (Chesnais et al., 2022) and cultured *in vitro* have spatially heterogeneous levels of NSP.

Here we confirm that NSH is a feature of endothelia *in vivo*. Our results using en face staining of murine aortas reveal heterogeneous levels of the transcription factor HES1 *in vivo* following similar patterns to those observed *in vitro*. These results highlight for the first time HES1 spatial heterogeneity in endothelia *in vivo*. Comparative analysis of our *in vitro* and *in vivo* data (Fig. 1B) confirms that NSH is preserved *in vitro* despite absence of concurrent signalling by perivascular cells and therefore enable us to use controlled *in vitro* experiments to unravel the causes of such heterogeneity.

It is challenging to measure the dynamics of endogenous NSP in human EC *in vitro* due to the absence of suitable gene promoter tracking systems. We have previously established the ECPT to enable high content single cell analyses of EC monolayers including spatial information. To understand whether the dynamics of NSP in EC is sufficient to generate heterogeneous levels of NSP downstream target genes, we present here a computational framework to evaluate the underlying molecular dynamics of NSP.

Many details of NSP dynamics have been previously elucidated using a host of experimental and computational methods (Boareto et al., 2015; Nandagopal et al., 2019; Sprinzak et al., 2010) allowing us to build a (parsimonious) set of possible and previously validated hypotheses. Overall, the dynamics of NOTCH signalling (excluding concurrent regulation by other signalling pathways) can be represented by five potential mechanisms: Lateral inhibition (Lib), lateral induction (Lid), productive or inhibitory interactions in cis (CI) and negative TF autoregulation (Fig. 2A)(Hirata et al., 2002).

To practically test alternative hypotheses, we encapsulated these mechanisms into a deterministic SMSM. Preliminary exploration of our SMSM revealed potential qualitative responses of NSP including emergent cell-contact mediated oscillations in TF (HES1) levels (Fig. 3 and SFig. 1). Oscillations in NOTCH downstream genes have been implicated and demonstrated in several biological context including somitogenesis and sensory organs development (Ferjentsik et al., 2009; Jiang et al., 2000). However, the standing hypothesis in the field is that such oscillations are associated with autoregulation of downstream TF such as HES1 rather than cell contact dependent mechanisms (Sturrock et al., 2011). Our SMSM reproduced very closely all other previously reported results including emergence of stable extreme and intermediate phenotypes (Boareto et al., 2015) and short-lived oscillation due to TF autoregulation (Fig. 3B). Nonetheless, we could not find parameters combinations for our SMSM which were able to generate self-sustained TF oscillations via TF autoregulation.

We reasoned that an EC specific transcription profile could affect the pace of NSP transduction which was previously estimated using transgene-mediated expression of ligands, receptors, and pathway reporters in NOTCH-inactive cell lines (LeBon et al., 2014; Nandagopal et al., 2019, 2018; Sprinzak et al., 2010). Time course experiments of EC exposed to the NOTCH inhibitor DAPT allowed us to estimate the characteristic timescale of signal transduction operating in different EC which are encapsulated by production and degradation parameters of key components in our ODE system. Overall, our experimental and in silico data closely aligned with previous estimates (Monk, 2003) providing further validation for our SMSM. Interestingly, results with HAoEC (Fig. 2C) points out at additional regulatory mechanisms which might encompass higher basal signalling in HAoEC as previously shown (Chesnais et al., 2022), which would lead to lesser sensitivity to DAPT at the concentration used in our experiment (and thus delayed inhibition). However, NSP-independent HES1 transcription has also been demonstrated (Curry et al., 2006) and it is likely to contribute to the net effects observed. Our experimental data show that both hypotheses are probably acting in concert as we can observe many EC with high HES1 intensity in both HUVEC and HAoEC monolayers upon long exposure with DAPT supporting the concept of NSP-independent HES1 transcription. At the same time, the NICD response in HUVEC was faster than in HAoEC (Fig. 2C), as NICD cleavage is upstream and independent from HES1, the simplest hypothesis to justify this effect is more abundant intrinsic NICD production in HAoEC as previously shown (Chesnais et al., 2022).

After calibration and validation of our SMSM we proceeded to evaluate whether any of the hypotheses was able to reproduce experimental data. To estimate fitting to experimental data we employed the Kolmogorov–Smirnov statistics. We used distribution of signal intensities in a cell map to build the empirical cumulative distribution functions (ECDF). The KS test was useful to exclude unfit hypotheses allowing us to test sufficiency of our SMSM to reproduce experimental data. After performing an exhaustive fine-grained parameter scan using an adaptive random search algorithm, we found several scenarios which were closely reproducing experimental data (Fig. 4A, KS D<0.06). Interestingly, all *in silico* results pointed at the importance of CI whereby all positive hits had cr values (affecting potency of CI) in the ~0.06-0.1 range corresponding to sustained CI. Importantly, the value of delay parameter involved in TF autoregulation (HES1 in our case) didn’t show similar weight in determining data fitting although some top positive hits (KS D<0.05) had delay values corresponding to ~15 min as previously estimated (Sturrock et al., 2011).

Detailed analysis of positive hits confirmed that these invariably corresponded to simulations predicting stable, periodic and asynchronous oscillations in HES1 and reproduced several previously reported experimental data including period of oscillations (~4-6 h) (Ubezio et al., 2016) and delay between NICD and HES 1 expression of ~20 minutes (Fig. 4B-D)(Curry et al., 2006).

We cannot rule out that cell-autonomous mechanisms can underpin HES1 oscillations only by analysis our SMSM because we implemented simple HES1 autoregulation as a sole hypothesis while further regulation layers not captured by our SMSM might be involved. For example, it is established that NSP oscillations during somitogenesis are driven by WNT and FGF signalling pathways (Carrieri and Dale, 2016). We therefore attempted to answer the question experimentally. We reasoned that, in the case cell-autonomous mechanisms underpin HES1 oscillations in EC we should observe heterogeneous levels of HES1 in cells cultured as sparse cells (absence of cell-cell contact). Our experiments using sparse EC cultures demonstrated that heterogeneity in NSP levels is lost under these conditions (Fig. 4E-F) and therefore strongly supports the hypothesis that oscillations are dependent on cell-cell interactions. These experiments also showed that NICD is actively transduced in absence of cell-cell contact confirming that CI in the same cell are sufficient to transduce NSP.

In the absence of other mechanisms, CI-mediated HES1 asynchronous oscillations are necessary and sufficient to generate spatially heterogeneous patterns of HES1 in simulated EC monolayer. Attempting extension to experimental data, we cannot rule-out different mechanisms than that proposed here to justify heterogeneity in NOTCH signalling. In fact, even our specific implementation of CI might represent a bias towards a specific but possibly not sufficiently general hypothesis. However, within these limitations our proposed mechanism is the most parsimonious model currently able to justify ours and other’s data regarding NSH in EC monolayers and warranting further experimental investigations.

Firstly, our results further support the idea that EC in the same monolayer acquire differential phenotypes and that these can be used by homogeneous cells (in terms of lineage) to exert differential functions in a highly parallel fashion. In this sense, emergent phenotypes and local EC connectivity have been already associated with endothelial plasticity, diversity of functions and robustness to perturbations (Chesnais et al., 2022; Lee et al., 2022; McCarron et al., 2019).

Secondly, our results strongly suggest that EC phenotypes can be modulated dynamically in timeframes of hours. This implies that each individual EC could in principle exert different sensor (ions, hormones, exogenous molecules), actuator (immune cells adhesion, solute trafficking), or maintenance (proliferation) functions at different moments in time while the overall balance of possible responses across the whole endothelium remains constant. We argue that implementing such heterogeneity dynamically (as we suggest) rather than with stable (fixed) phenotypes renders the endothelium more robust to perturbations. For example, if specific EC subtypes are lost due to a pathogenic insult the remaining cells could easily replace the lost ones by proliferation without loss of function and without need for specific EC precursors/stem (Chesnais et al., 2022; McCarron et al., 2019).

Finally, our data strongly suggests that the dynamics of NOTCH signalling in EC contains non-linearities dependent on molecular, geometrical, and spatial constraints which introduce challenges in predicting these dynamics in living organism and thus understanding the dose-effects responses of pathway modulator drugs.

In conclusion, in the present work we offer a framework for the default state of EC in healthy monolayers. A vast amount of research is ongoing to understand and exploit molecular mechanisms underpinning EC functions in the context of physiologic (wound healing), defective (diabetes) or pathologic (cancer) angiogenesis characterised by rapid changes in spatial relations between EC. The consequences of the hypotheses raised and developed have implications in designing how to perturb these mechanisms to treat diseases.

## Materials and Methods

### Cell culture

All *in vitro* data shown in the present work has been generated as described previously (Chesnais et al., 2022). Briefly, HAoECs and HUVECs (PromoCell) were plated on 10 μg/ml fibronectin (Promocell)-coated flasks, grown in EGMV2 medium (Promocell), detached with accutase (Thermo Fisher Scientific, Waltham, MA), and used by passage 5. For experiments, 4×10^4^ ECs per well were seeded for confluent experiments and 1×10^3^ ECs per well for sparse experiments in fibronectin-coated 96-well plates (μClear, Greiner). Cells were cultured for 24h for sparse and 96h for confluent under basal conditions (EGMV2, Promocell) or treated with DAPT (5μM, Tocris bioscience, UK).

### Immunostaining and image acquisition

Cells were fixed with 2% paraformaldehyde in phosphate-buffered saline (PBS) for 10 min at room temperature. Cells were blocked 1 h with PBS supplemented with 1% fetal bovine serum (FBS) and permeabilised with 0.1% Triton X-100. Cells were then incubated for 1 h at room temperature with primary antibodies against CDH5 (VE-cadherin; Novusbio NB600-1409, 1μg/ml final), NOTCH1 (Abcam, ab194122, Alexa Fluor 647-conjugated, 1 μg/ml final) and Hes1 (Abcam, ab119776, 1 μg/ml final). Plates were washed and incubated 1 h with 1 μg/ml secondary Alexa Fluor 488-conjugated and Alexa Fluor 555-conjugated antibody (Thermo Fisher Scientific), Hoechst 33342 (1 μg/ml, Sigma). We obtained the images with an Operetta CLS system (PerkinElmer, Waltham, MA) equipped with a 40× water immersion lens (NA 1.1).

### En face preparation and whole mount staining of mouse aorta

For en face preparation of aortas, 2–8-month-old mice (C57BL/6J, Jax Strain 000664) euthanised for standard colony maintenance under Schedule 1 methods were used. For preparation and immunostaining, we used an adaptation of a previously established protocol (Hakanpaa et al., 2015). In brief, upon euthanasia animals were immediately perfused with PBS by intracardial injection and subsequently perfusion fixed by injection of 2% PFA–PBS. Dissected aortas were fixed for additional 1□h by immersion in 2% PFA–PBS, washed extensively with PBS and blocked with FBS (5% FBS + 0.01% Triton-X) overnight at 4°C. Immunostaining was performed after permeabilization (0.1% Triton-X for 10minutes at RT) using primary antibodies diluted in blocking solution (VE-Cadherin, #14-1441-82, 5 μg/ml final and HES1, #PA5-28802, 5 μg/ml final, Thermo Fisher Scientific) for 48h at 4°C, followed by extensive washing using blocking solution at RT. Aortas were then incubated with Alexa Fluor 488-conjugated and Alexa Fluor 555-conjugated antibody (Thermo Fisher Scientific) and Hoechst 33342, for 24h at 4°C, washed and mounted on a coverslip with Mowiol (Sigma-Aldrich). The aortas were imaged, and z-stacks were obtained using a Leica Sp8 confocal microscope with a 40X air objective.

### Image and data analysis

Analysis of images obtained in this study has been done with ECPT (endothelial cells profiling tool) (Chesnais et al., 2022).

### Spatial model of cell monolayers in Compucell 3D

The multicellular spatialised model of EC monolayers was developed using a cellular Potts model formalism (CPM, or Glazier-Graner-Hogeweg model (Swat et al., 2012) using the software Compucell3D (CC3D version 4.2.5, www.compucell3d.org,). Cell maps were imported as .piff files.

### Generation of cell maps

To generate cell maps extracted from in vitro experiments we used our high content image analysis platform (Chesnais et al., 2022) and developed custom imageJ and R scripts to generate .piff files which can be inputted in CC3D.

### CC3D model modules

We implemented a CC3D model including five “steppables”: Input/output, SBML solver, Visualisation, Cell Initialisation and Delta/Notch neighbour interactions. The Antimony code encoding the ODE model and call to all steppables are included as additional modules. The first four steppables are standard CC3D modules and extensive documentation on how to setup such modules is available on the CC3D website and manuals. The fifth stoppable, NDJ_Interactions, is dedicated to evaluating neighbours’ interactions as described in results. The module calculates contact areas (CA) between each cell and its neighbour’s CA is then factored when calculating amount of trans interactions between individual pairs of cells at each MCS thus linking cell shape and disposition with extent of NOTCH signalling.

### Ordinary Differential Equations model of NOTCH signalling

We developed the mathematical framework of NOTCH signalling as an ordinary differential equations (ODE) system as proposed previously (Boareto et al., 2015; Sprinzak et al., 2010) and encoding a hybrid protein/gene regulatory network (P/GRN) using the Tellurium package and the Antimony language (SBML compatible). We assumed a well-stirred bioreactor cell model where we explicitly encoded and carefully calibrated time delays in selected reactions and otherwise assumed that reaction steady state would be reached in a time scale smaller than our individual MCS (5 minutes).

We firstly drafted laws for production of Dll4, Notch1, Jag1 and Notch4 and downstream target genes Hes1 and Hey1/2 which can modulate production of the above.

For all production laws we used Hill functions which have been previously shown to recapitulate the dynamics of TF mediated gene expression and protein production in the NOTCH pathway (Boareto et al., 2015; Sprinzak et al., 2010):

For positive regulation (Hp):

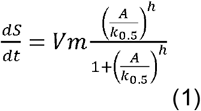

For negative regulation (Hn):

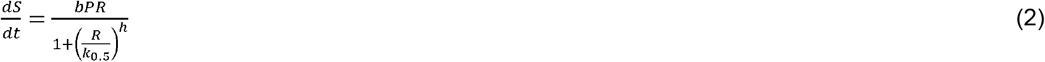

For competitive regulation (Hc):

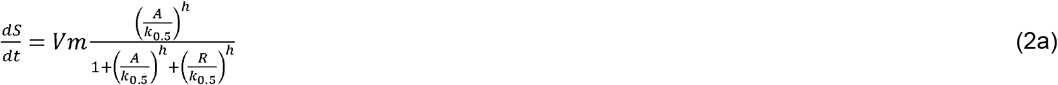

Were *Vm* is the maximal production rate of the molecule *S, bPR* represent the basal production rate, *A or R* represents the concentration of regulatory transcription factor (A for Activator, R for Repressor), *k_0.5_* represents the concentration of TF exerting half maximal effect and *h* is the Hill coefficient widely used to represent cooperativity and dimerization processes (HES1 functions as homo/hetero-dimer) and previously shown to appropriately reproduce the mechanisms discussed in this work (Boareto et al., 2015). For degradation laws and productive/non-productive cis interactions we used mass action functions.

Mass action (Ma):

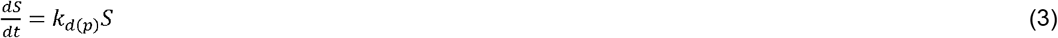

Where *k_d_* or alternatively *k_p_* represent degradation or production rates respectively.

The full model for species production is as follows:

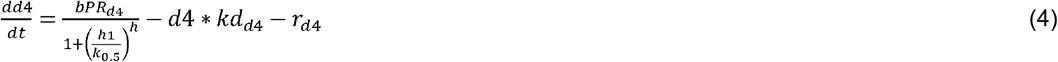

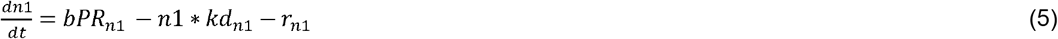

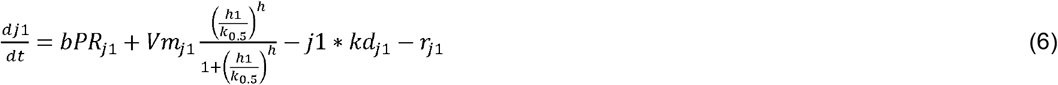

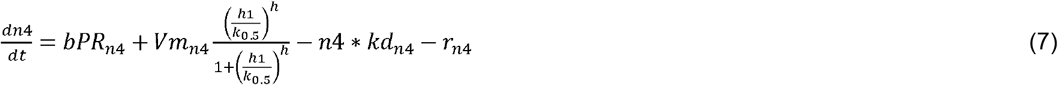

HES 1 production in absence of autoregulatory feedback:

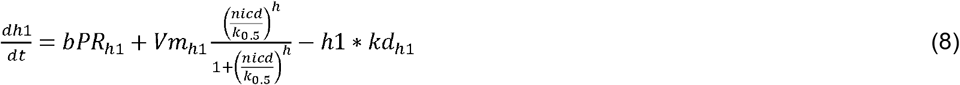

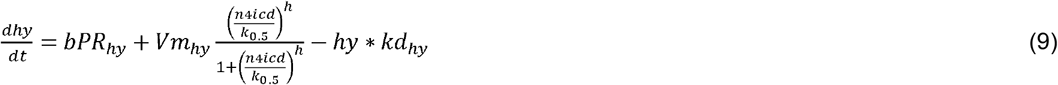

Where *rd4* and *rj1* are the amount of “Reacted” ligand which is consumed in cis and trans dimerization events leading to the production of cleaved Notch intracellular domains, the second messengers of Notch signal transduction. *Rd4* and *rj1* are calculated in the CC3D environment while *rn1* and *rn4* are calculated via a Hill function parallel to Eqs 10 and 11 below *(Hp of tmdn or tmjn respectively)*. All the other species abbreviations as per Table 1.

### NICD production

Eq 8 and 9 (Hes1 and Hey1/2 production) depends on the two species *NICD* and *N4ICD* which represent the second messengers (notch intracellular domain, NICD) of Notch1 and Notch4 respectively. Production laws for *NICD* and *N4ICD* use transit parameters as inputs (*tmDN, tmJN*). *tmDN* and *tmJN* are calculated in the CC3D environment from the interactions in cis and trans of relevant ligands and factoring differential adhesion between neighbouring cells. The laws governing *tmDN* and *tmJN* calculations are discussed below. This is the key interaction point between CC3D python and Antimony/SBML codes allowing to account for spatial disposition and differential contact between cells. *NICD* and *N4ICD* production laws are shown in eq 10 and 11.

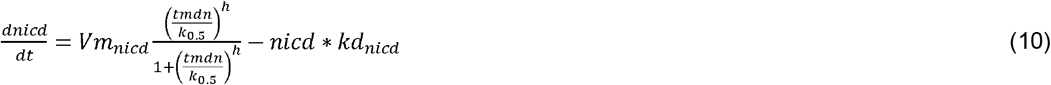

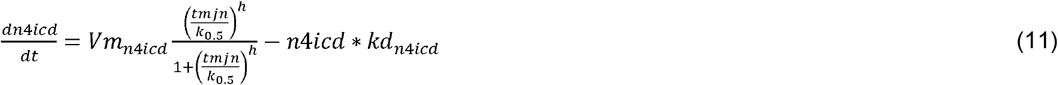

### Cis interactions

Previous models (Boareto et al., 2015) have only accounted for cis-inhibitory effects whereby membrane interactions of Notch ligands with receptors lead to dampening of the signalling by limiting ligands available for productive trans interactions. In our model we also included productive cis interactions which have been recently demonstrated (Nandagopal et al., 2019) and supported by our own data (Isolated cells).

Non-productive competitive cis- interaction (cJN) between Jagged1 and Notch1 follows a mass action law limited by the lesser abundant of the two proteins as follows:

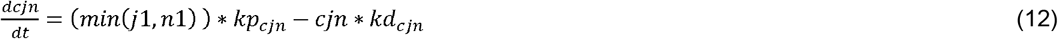

Where *min(j1, n1)* is the minimum value among *j1 and n1* in the same cell.

Productive cis interactions (*cdn*) between Dll4 and Notch1 follows a similar law but differently from *cjn, cdn*, it can be converted in an active specie (*cadn*) participating in NICD production as follows:

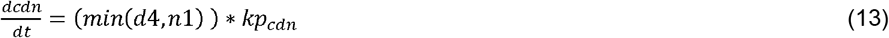

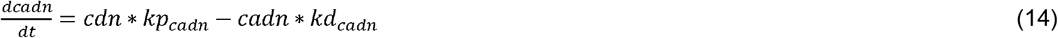

The law for NICD production including productive cis interactions becomes:

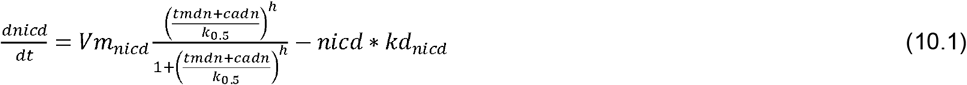

The *k_p_* parameter for production of *cjn* and *cdn* is not fixed but dependent from a scaling parameter inversely proportional to trans interaction (*tmdn, tmjn*) to capture a reported concept whether strength of cis interaction is inversely proportional to strength of cell adhesion and consequent trans interactions (Zhou et al., 2022).

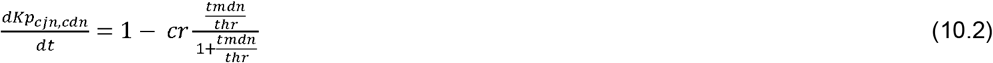

### HES1 autoregulation

We implemented the reported HES1 autoregulatory mechanism in our model as a competition mechanism between NICD and HES1. To encode delay, required to produce oscillations according to previous reports, we explicitly included an intermediate species in the HES1 processing chain. No intermediate factor has been reported in the literature so far, therefore we assumed that the delay is caused by nuclear import/export of involved species as previously suggested. The production law for HES1 including autoregulation becomes

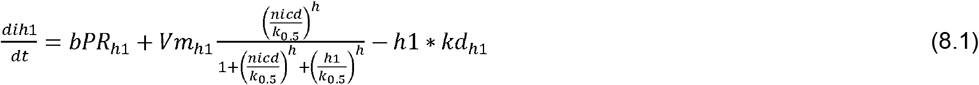

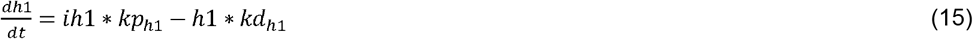

Where *ih1* (inactive HES1) represent inactive species such as mRNA and cytoplasmic protein before nuclear import. Therefore, active h1 production is delayed by a scaling factor (*kp_h1_*) in the first term of 15.

The full Antimony version of the model is included in supplementary material and provided as a working Tellurium script. The Tellurium script can be used to test a single cell model where *tmDN, tmJN, Rd4, Rj1* (and all the other parameters in the simulations can be modulated. The model as described has steady states solutions for all parameter ranges used in this study (SFig. 1B). Full parameter list and corresponding values are provided in table 1.

### Calculation of trans interactions (DeltaNotchNeighborSteppable)

As discussed above, *tmDN* and *tmJN* (and corresponding *Rd4* and *Rj1* in eq 4 and 6) parameters are calculated in the CC3D python environment and then inputted into the Antimony/SBML model at each MCS. In brief, *tmDN* and *tmJN* are calculated by evaluating amount of contact between each cell and each of its neighbours and trading the relevant ligands. To account for competition between Dll4 and Jagged1 for Notch1 receptor we imposed a partitioning parameter (kpDJ, calculated each MCS) which is proportional to Dll4 concentration and inversely proportional to Jagged 1 concentration. It is established that also the strength of Jagged1 affinity for Notch1 is modulated dynamically by posttranslational modifications (Zhou et al., 2022) and to capture this possibility we imposed a further weighting parameter (KJC). However, analysis of this level of Notch signalling modulation was outside the scope of the present work and the KJC parameter has been fixed to 1 (equal affinities of Dll4 and Jagged 1) in all our simulations shown in results.

### Statistical analysis

All data wrangling and statistical analyses were performed in R Studio using the Tidyverse package except the KS test performed runtime for random sampling experiments as discussed below.

To implement a random search algorithm searching parameter sets fitting our experimental data we firstly modified our model to work as a black-box function accepting parameters sets as inputs and returning a metrics of data fit as output. As data fit metrics we used the KS distance as implemented in the numpy library, experimental data were randomly sampled from our database in R and stored in a file accessible to the simulation while synthetic ones were collected runtime. To run the random search, we developed a Python script which randomly sampled the parameter space with a latin hypercube sampling strategy, instantiated individual simulations for each sample, collected and compared results and then iteratively optimised the results according to the schematics in SFig. 3. We repeated the procedure 100 times for a total of >30000 simulations. Average runtime of all simulation was ~20s/simulation (each simulation running for 150 MCS ~ 15 hours) on a Mac Book Pro 2019 (8-cores Intel i9, 2.3GhZ, 64Mb RAM).

## Acknowledgments

The authors wish to thank Prof Fiona Watt for giving access to mouse tissue and facilities for tissue microdissection and imaging.

---

**Supplementary Fig 1:**
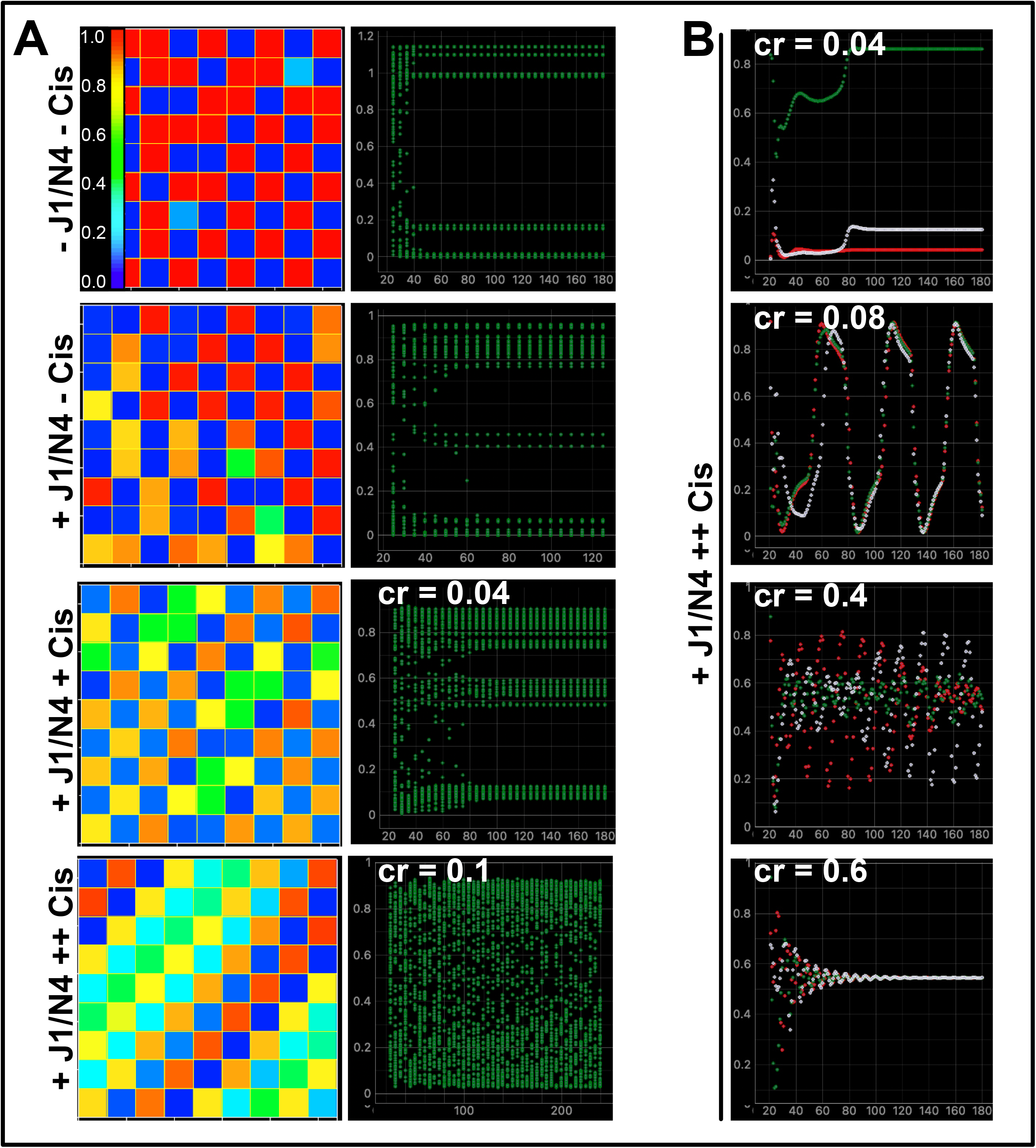
SMSM validation with regularly shaped and distributed cells. **A)** Representative maps (left panels) and timeseries for all cells (right panel) for the indicated conditions (−= OFF, += ON, cr=0 or as indicated in individual panels). **B)** Time series traces for 3 random cells illustrating the qualitative effect of increasing cr, Cr values>0.1 were not considered biologically plausible considering the other parameters in our SMSM and corresponding simulations were not analysed further in the present work.

**Supplementary Fig 2.:**
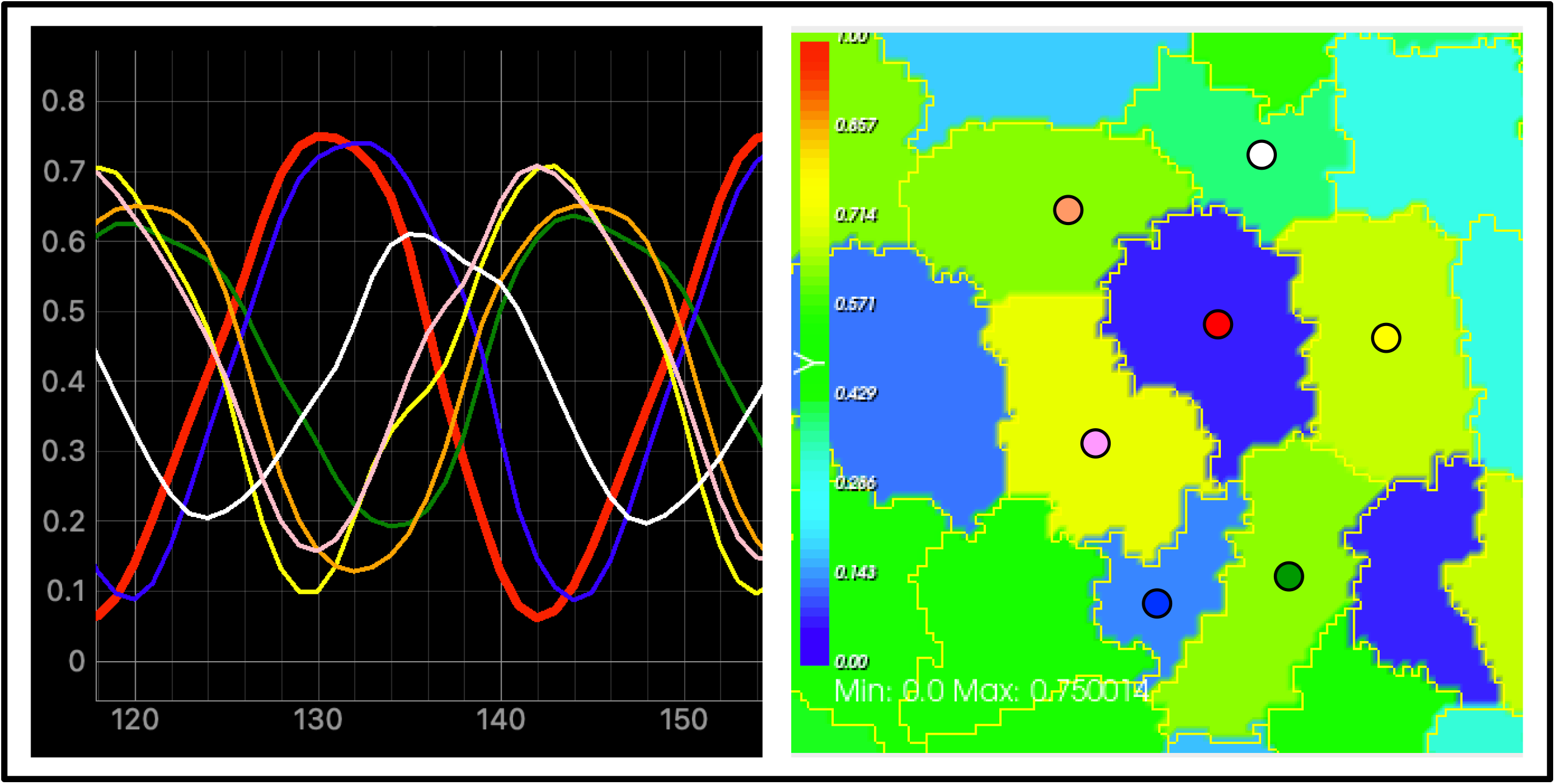
Periodic oscillations in cell neighbourhoods are asynchronous. Representative time series (left panel) and corresponding map (right panel) of NSP signalling in a neighbourhood of six cells under the parameter setup of Fig 4B. Colour of dots in right panels correspond to colour of traces in left panel. The red trace corresponds to the central cell in the neighbourhood. In a typical scenario, the signal in different neighbouring cells can be synchronised to the central cell in phase (blue trace) or anti-phase (yellow, pink traces). However, signal dynamics is typically asynchronous in several cells (orange, white and green traces). Scenarios encompassing full synchronisation with central cell either in phase or anti-phase are not observed under physiologically relevant parameter setups.

**Supplementary Fig 3.**
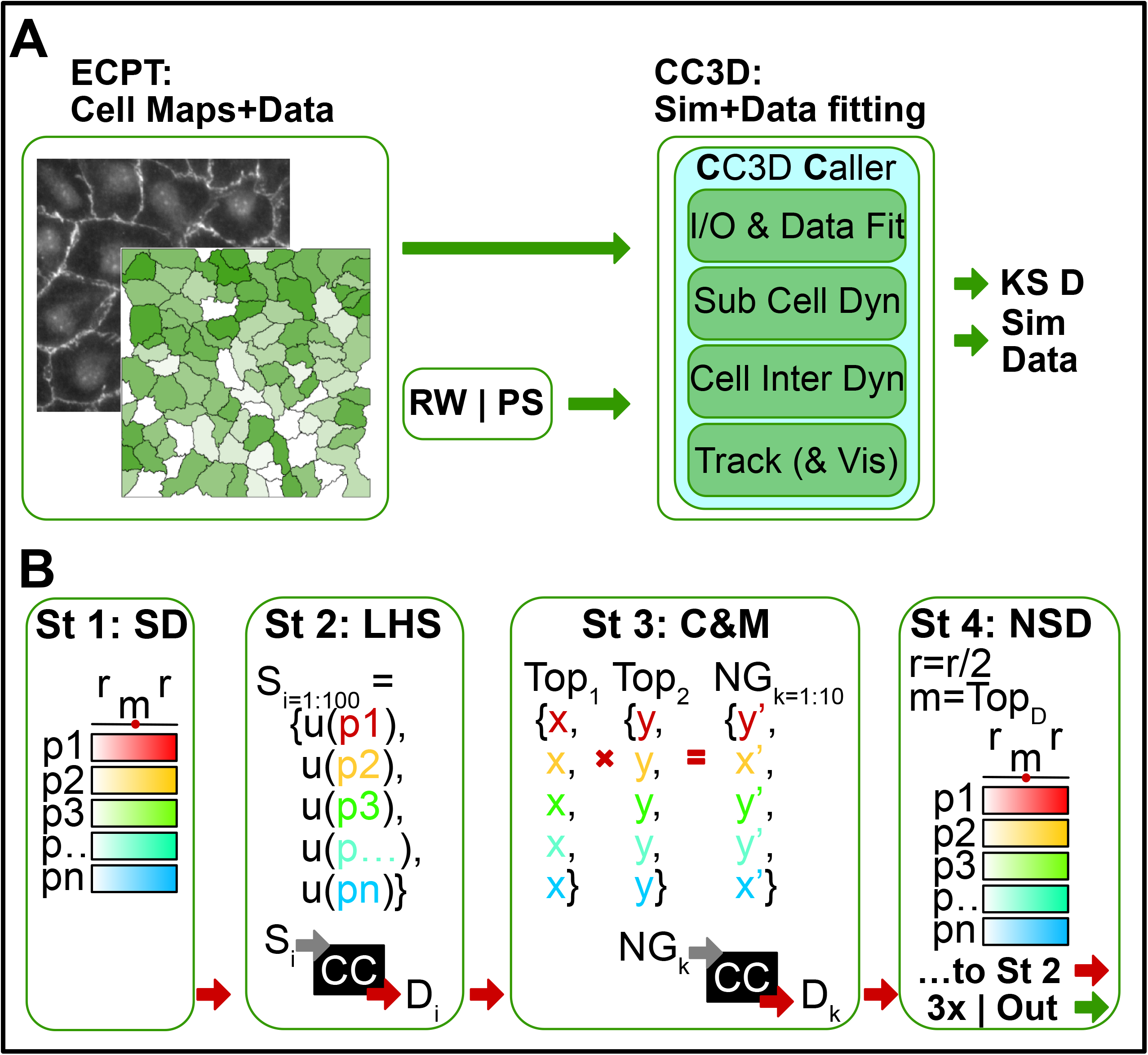
Random search strategy for fine-grain parameter scan. **A)** Schematics of the strategy employed in parameter scan. ECPT measurements and maps, and Random Walk (RW) or Parameter Scan (PS) inputs are fed to a black box CC3D caller. The black box includes all modules of the SMSM and one additional module performing KS test runtime. The output of this function is the KS distance and/or time series data corresponding to simulation parameters. **B)** Random walk sequence. St 1: Sample definitions. Mid value (m) and range (r) for the selected parameters are provided as user input. St 2: Latin Hypercube Sampling is performed on selected parameters to generate a sample set of desired size (i=100), Each sample is used as input for the CC3D Caller (CC) retuning a corresponding KS distance metrics between synthetic data and reference data (D_i_ ). St 3: Crossing and mutation. Top 2 results from St 2 are selected and their input parameters (for example, x for T1 and y for T2) crossed k times to produce a new generation (NG) with k elements. X or y are assigned randomly to each KG_k_ and they are also mutated to an user defined extent (r/5 for results shown in this manuscript). Individual simulations are then run for each individual element of the new generation. St 4: The top result from St 3 (either T1 or NG_k_) is used as basis to generate a more focussed sample definition (m = Top_D_, r=r/2) and inputted as new iteration to St 2. Iterations number (x) and other parameters are all user defined and we used values indicated in figure for the results shown in the manuscript.

## Notes

### Competing Interest Statement

The authors have declared no competing interest.

