## Supplementary material for "A spatialised agent-based model of NOTCH signalling pathway in Endothelial Cells predicts emergent heterogeneity due to continual dynamic phenotypic adjustments": Table 1

| | UniprotKB | Vm | bPR | h | k50 | Kd | Model Eq N | ~typical mol/cell (bPR = 1) |
|---|---|---|---|---|---|---|---|---|
| NOTCH1 | P46531 | 1 | 0.6-1.2 | 4 | 50 | 1 | 5 | 2500 |
| DLL4 | Q9NR61 | 1 | 0.6-1.2 | 4 | 50 | 1 | 4 | 2500 |
| NOTCH4 | Q99466 | 0.0-1.0 | 0 | 4 | 50 | 1 | 7 | 2500 |
| JAGGED1 | P78504 | 0.0-1.0 | 0 | 4 | 50 | 1 | 6 | 2500 |
| HES1 (cytopl) | Q14469 | 1 | 0 | 2 | 50 | - | 8-8.1 | 0-100 |
| aHES1 (nucl) | Q14469 | 0.01-1 | 0 | - | - | 0.09 | 15 | 0-100 |
| HEY2 | Q9Y5J3 | 1 | 0 | 2 | 50 | 0.09 | 9 | 0-100 |
| cDN | - | Kpcdn | - | - | - | 0.1 | 10.2-13 | - |
| caDN | P46531 | 0.1 | - | - | - | 0.05 | 14 | - |
| cJN | - | Kpcjn | - | - | - | 0.1 | 10.2-12 | - |
| NICD | P46531 | 100 | 0 | 4 | 2000 | 1 | 10-10.1 | 0-100 |
| N4ICD | Q99466 | 100 | 0 | 4 | 2000 | 1 | 11 | 0-100 |
| cr | 0.0-0.1 | | | | | | | |
| KJC | 1 | Weighting of J1/D4 competition for N1 | | | | | | |

Table 1 Parameters values used in the NSP SMSM. Scanned parameters highlighted in green.
